# MatriSpace: Identification and visualization of spatially resolved ECM gene expression patterns in health and disease

**DOI:** 10.64898/2026.04.26.720198

**Authors:** Ayomide Oshinjo, Daiqing Chen, Petar Petrov, Valerio Izzi, Alexandra Naba

## Abstract

The extracellular matrix (ECM) is a highly dynamic network of proteins forming the structural organizer of all tissues. Different cell populations contribute to the assembly of the 150+ proteins of a functional ECM. In addition, different ECM subtypes, supporting distinct cellular functions, are found in every organ. Spatial transcriptomics (ST) provides a unique, yet untapped, opportunity to identify which cell populations contribute to ECM production with spatial context. Applied to healthy and diseased samples, this method can identify ECM changes that could be exploited for therapeutic purposes. Here, we introduce MatriSpace, a computational framework to mine ST datasets with a focus on ECM genes. MatriSpace offers two operating modes: researchers can either upload their own ST datasets or explore a large collection of public datasets. Upon analysis, MatriSpace returns spatially resolved maps of matrisome gene expression in relation to cell populations, at multiple levels: from single-gene analysis to tissue niches and functional ECM units. MatriSpace is available as an R package and an online Shiny App (https://matrinet.shinyapps.io/matrispace), making it accessible to all users regardless of their level of expertise.

**Graphical abstract:** 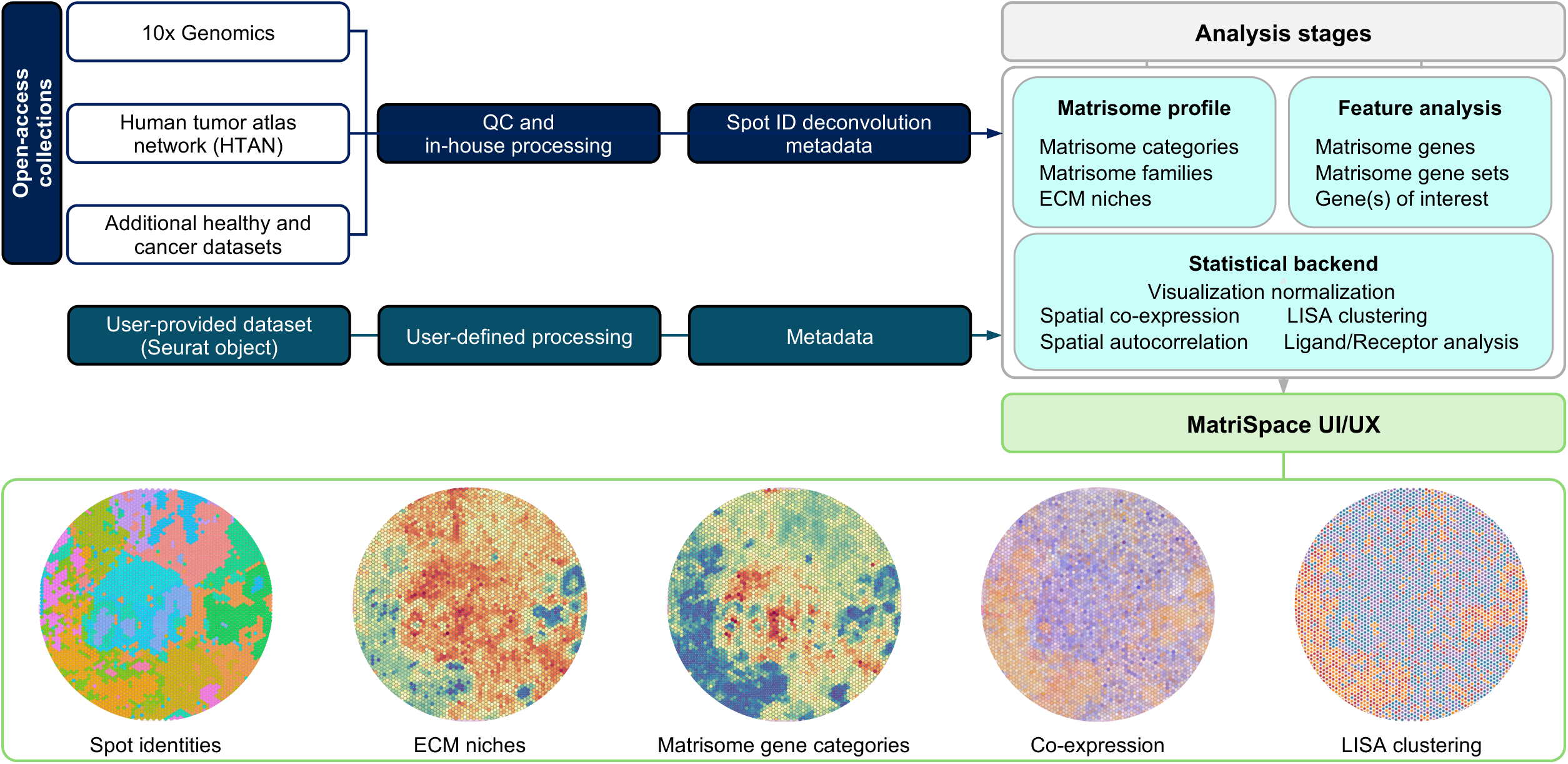

**KEY POINTS:**

- MatriSpace is a computational framework to interrogate ECM gene expression in spatial transcriptomic datasets.
- Researchers can upload their own spatial transcriptomic datasets for processing by MatriSpace.
- Researchers can interrogate a vast collection of public datasets of healthy and diseased tissues through MatriSpace.
- MatriSpace can identify, quantify, and interpret spatial expression patterns of matrisome genes, gene sets, and niches.
- MatriSpace can uncover regional coordinations of matrisome components and their relationships with non-matrisome genes, such as matrisome receptors.

## INTRODUCTION

Spatial transcriptomics (ST) is a powerful approach that enables simultaneous measurement of gene expression and spatial context within intact tissues. ST data can be leveraged to infer the distribution of cell populations and subpopulations within tissues and identify neighborhood interactions, cell lineages, signaling niches, and emergent tissue architectures, such as gradients of hypoxia, immune infiltration, or stromal activation (Honcharuk et al., 2025; Moses and Pachter, 2022; Rao et al., 2021; Ståhl et al., 2016). ST is thus particularly well-suited to study inherently spatial systems.

The extracellular matrix (ECM) is a complex network of proteins that governs cell organization and tissue architecture. It is the product of the coordinated production, secretion, and assembly of proteins, governed by tightly controlled gene expression programs, across multiple cell populations. Two main subtypes of ECMs, the basement membrane ECM and the interstitial ECM, characterized by different molecular compositions, functions, and cellular interactions, coexist in tissues and organs (Naba, 2024). In addition, the ECM undergoes constant remodeling during developmental processes, in homeostatic tissues, but also in pathological processes such as tissue repair, cancer, and fibrosis (Bonnans et al., 2014). Single-cell RNA sequencing has revealed that distinct cell subpopulations contribute to the production of enzymes that remodel the ECM. Different cell populations also express different subsets of ECM ligands and ECM receptors, accounting for the autocrine and paracrine signaling functions of the ECM (Lamba et al., 2025). The ECM is thus a remarkable example of a spatial system, with locoregional specification, that has direct functional consequences.

It is thus evident that ST can be leveraged to map ECM-gene-expressing cell populations (*e.g.*, fibroblast subtypes), resolve spatial patterns of matrisome gene expression, identify ECM-rich niches such as fibrotic or desmoplastic regions, and infer how ECM composition correlates with complex tissue-wide processes, such as immune infiltration or exclusion, tumor progression, or tissue regeneration. Current analytical frameworks for ST, however, are poorly suited to capture the complexity of ECM biology. Most pipelines are optimized for cell-centric questions such as clustering, cell-type annotation, and ligand–receptor inference (Kalantari-Dehaghi et al., 2025).

Since most proteins are very long-lived, ECM gene expression levels are typically low, especially in homeostatic tissues, and, as discussed, ECM genes are typically expressed by multiple cell types, making their analysis difficult to resolve with standard normalization, dimensionality reduction, and clustering approaches. Additionally, existing tools typically ignore key ECM properties such as supramolecular assembly and long-range continuity, which are not encoded in single transcript counts but rather emerge from spatial coordination of gene expression programs. As a result, current ST analysis platforms fail to capture the mesoscale organization of ECM networks and allow only exploratory, unsystematic analyses of the matrisome. For example, Pentimalli and colleagues used ST to profile cell types and map cell-ECM interactions in 3D in lung carcinoma samples (Pentimalli et al., 2025). De and colleagues employed ST to characterize the molecular composition of glioblastomas. They identified that different subsets of glioblastoma cells, found in distinct tumor niches, express distinct ECM gene signatures and engage in different signaling networks (De et al., 2025). High-resolution mapping of the tumor microenvironment using integrated single-cell, spatial, and *in-situ* analyses further shows how ECM features co-vary with local cell states in breast cancer (Janesick et al., 2023) and melanoma (Hunter et al., 2021). Together, these works provide important descriptive insights into ECM spatial specialization, yet lack a systematic, quantitative framework for defining ECM-driven functional tissue units. These limitations, together with the rapid adoption of ST by the scientific community, motivated the development of MatriSpace, a platform specifically designed to account for the specificity of the ECM and treat it as a spatially continuous, integrated system. MatriSpace offers two operating modes: researchers can either upload their own datasets or query a collection of nearly 200 public datasets. Upon analysis, MatriSpace returns spatially resolved maps of matrisome gene expression in relation to cell populations, at multiple levels: from single-gene analysis to tissue niches and functional ECM units. MatriSpace is available as an online Shiny App (https://matrinet.shinyapps.io/matrispace) and as an R package (https://github.com/izzilab/matrispace), making it accessible to all users, regardless of their level of expertise in bioinformatics. Through two case studies, we further illustrate how, beyond providing a molecular readout, MatriSpace can help gain insights into ECM function and, hence, tissue function.

## METHODS

### Overview of MatriSpace architecture

MatriSpace is a computational toolbox for analyzing matrisome gene expression in spatially resolved transcriptomics datasets. It is distributed through three complementary components tailored to different user needs. All components rely on a shared analytical backend implemented in R Statistical Software v4.3.3 (R Core Team, 2024), ensuring full reproducibility across deployment modes. Access to MatriSpace is offered via:

– An online web application freely accessible at https://matrinet.shinyapps.io/matrispace, which provides access to a collection of pre-processed 10x Genomics Visium datasets sourced from public repositories, in addition to user uploads of up to 1 GB.
– An offline application, available at https://github.com/izzilab/matrispace, is accessible to users who prefer local computation with the ease of use of the MatriSpace user interface (UI). This option removes upload size limitations but does not include the preloaded dataset collection.
– Last, the MatriSpace R package, available at https://github.com/izzilab/matrispace, provides programmatic access to all analytical functions and supports seamless integration into Seurat-based pipelines.

### Data input

#### Sourcing of open-access sample datasets

The preloaded dataset collection comprises 198 10x Genomics Visium datasets obtained from public repositories, including 10x Genomics (https://www.10xgenomics.com/datasets), the National Cancer Institute’s Human Tumor Atlas Network (https://humantumoratlas.org/; (Rozenblatt-Rosen et al., 2020)), the Gene Expression Omnibus (GEO, http://www.ncbi.nlm.nih.gov/geo; (Barrett et al., 2013)), and Zenodo (https://zenodo.org/) (**Supplemental Table S1**). The open-access collection is divided into healthy datasets (18) and cancer datasets (180). To ensure analytical robustness, all samples underwent stringent quality control using the standard Seurat SRT preprocessing workflow. Datasets with fewer than 500 detected genes, fewer than 100 tissue-covered spots, or poor tissue section quality were excluded.

#### User data upload and processing

MatriSpace supports user data input as Seurat objects (.rds) or SpatialExperiment objects (.rds), ensuring compatibility with both the Seurat and Bioconductor ecosystems. SpatialExperiment objects are automatically converted to Seurat format while preserving spatial coordinates, tissue images, and spot-level metadata. Upon upload, datasets are processed through an automated pipeline that standardizes gene symbols, applies variance-stabilizing normalization when required, computes matrisome gene set and niche scores, and annotates ECM niches. To avoid redundant computation, the pipeline automatically detects and retains any pre-existing user-defined annotations.

### Annotation options

All MatriSpace analyses are stratified by user-selected annotation variables. For the preloaded dataset collection, three default annotations are provided:

– Seurat clusters represent unsupervised groupings based on gene expression similarity.
– Major cell types define broad cellular identities, including fibroblasts, endothelial cells, and immune populations.
– Cell subtypes provide higher-resolution annotations, such as distinguishing regulatory from effector CD4^+^ T cells.

Annotation pipelines were adapted to account for biological differences between tissue types. Healthy tissue samples were annotated using the ScType marker-based classifier (Nader et al., 2024), with major cell types defined using the ScType reference database and cell subtypes derived from the CellMarker 2.0 database (Hu et al., 2023). Cancer samples were annotated using SpaCET, which estimates continuous cell-type proportions for each spot via deconvolution (Hao et al., 2024). These proportions are converted to categorical labels by assigning each spot to the cell type with the highest proportion, provided it exceeds a 10% threshold. SpaCET also allows mixed identities for spots containing multiple populations. For spots where malignant cells are the dominant population and where a second non-malignant cell type also exceeds a secondary threshold, a combined label is assigned (*e.g.*, “Malignant+Endothelial”). This threshold was selected to maximize spatial coverage, as each Visium spot (∼50 μm in diameter) typically captures 10 to 50 cells, making single-cell-type purity rare.

Independent of cellular identity, each spot is also classified as being part of an interstitial or basement membrane ECM niche using marker-based scoring (*see below*). Last, for user-uploaded data, any metadata column present in the object can be selected for downstream analyses.

### Analytical workflows

#### Matrisome profiling

The matrisome profiling workflow computes expression scores for a panel of gene sets designed to inform on the structure or function of the proteins they encode. These gene sets include: **1)** the canonical matrisome gene sets that classify core ECM and ECM-associated components in six gene sets based on the domain-based organization of the proteins they encode (Hynes and Naba, 2012; Naba et al., 2012). These include: ECM glycoproteins, collagens, proteoglycans, ECM-affiliated, ECM regulators, secreted factors (**Supplemental Table S2A**); **2)** four matrisome gene families: genes encoding laminins (Domogatskaya et al., 2012), matricellular proteins (Bornstein and Sage, 2002), and two classes of transmembrane proteoglycans, the glypicans and syndecans (Iozzo and Schaefer, 2015) (**Supplemental Table S2B**); **3)** four functional subcategories of matrisome genes, namely, genes encoding ECM proteins involved in hemostasis (Bergmeier and Hynes, 2012), genes encoding proteins found in the perivascular region, genes encoding matrisome proteins participating in the formation of elastic fibers (Baldwin et al., 2013), and genes encoding growth-factor-binding ECM proteins (Hynes, 2009) (**Supplemental Table S2C)**. Each spot is also assigned to one of two functional ECM niches, the basement membrane ECM (41 signature genes; (Page-McCaw and Ferrell, 2025; Yurchenco, 2011)) or the interstitial ECM (22 signature genes; (Hynes and Naba, 2012; Naba, 2024)), based on the expression of curated gene sets provided in **Supplemental Table S2D**. These gene sets were built based on prior knowledge and through the curatorial work of the Naba lab as part of the Gene Ontology consortium (https://geneontology.org/) (Gene Ontology Consortium, 2025).

#### Feature analysis

Users may interrogate specific features using a two-level selection system. The primary feature can be a matrisome gene or gene set, while the optional secondary feature (including additional subsets of matrisome genes; **Supplemental Table S2E**) may be any detected gene or matrisome gene set. The output of the feature analysis includes spatial feature maps with autocorrelation statistics, expression distributions across annotation groups, co-expression blend plots visualizing joint spatial patterns, and local spatial association maps classifying spots based on their spatial relationships (*see Results*).

### Statistics and data visualization

#### Matrisome gene set scoring

Matrisome gene set expression scores are computed using UCell, a rank-based method robust to differences in dataset composition (Andreatta and Carmona, 2021). For a gene expression matrix with *g* genes across *c* spots, UCell ranks genes within each spot. For a gene set of size *n*, the UCell score for spot *j* is defined as:

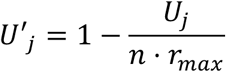

where *U* is the Mann-Whitney U statistic:

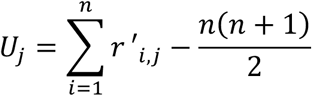

Scores depend solely on within-spot gene ranking, rendering them robust to batch effects and variability in capture efficiency.

#### ECM niche classification

ECM niche assignment is performed using a marker-based scoring strategy adapted from ScType (Nader et al., 2024). For each spot *i* and niche *d*, an enrichment score is computed as

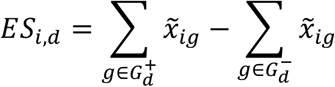

where 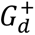 and 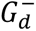 denote positive and negative marker gene sets for niche d, and *x̃_ig_* is the scaled expression of gene *g* in spot *i*. Each spot is assigned to the niche with the highest positive enrichment score, if any, or remains unclassified if all scores are uniformly negative.

#### Visualization normalization

Two complementary normalization schemes are applied for visualization. Robust normalization highlights baseline spatial patterns while limiting outlier influence:

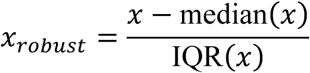

Log-scaled normalization is used for hotspot detection to emphasize focal regions of high expression:

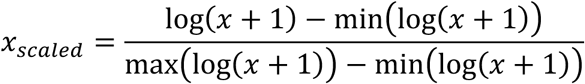

#### Spatial autocorrelation

Spatial organization of gene or gene set expression is assessed using Moran’s *I*, implemented via the MERINGUE package (Miller et al., 2021).

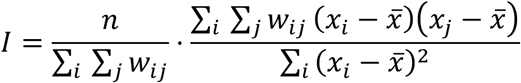

where *n* is the number of spots and *w_i,j_* represents inverse-distance spatial weights. Statistical significance is evaluated by permutation testing.

#### Feature expression and pairwise correlation

Differences in feature expression across annotation groups are assessed using one-way ANOVA. When two features are selected, their association is evaluated using both Pearson correlation.

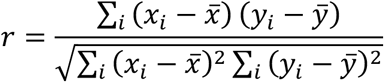

and spatial cross-correlation, which incorporates spatial context via inverse-distance weights (MERINGUE):

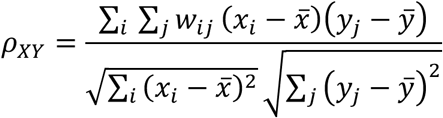

where *w_ij_* is the spatial weight between spots *i* and *j*. This dual approach distinguishes linear co-expression from spatially coordinated expression. Statistical significance is evaluated by permutation testing.

#### Co-expression blend visualization

Bivariate spatial expression is visualized using a four-corner color gradient. After normalization to a common scale, expression levels are mapped to a two-dimensional color space: white indicates low expression of both features, primary and secondary colors indicate dominance of individual features, and blended colors represent co-expression. This visualization enables rapid identification of co-localized, mutually exclusive, or jointly absent expression patterns.

#### Local indicators of spatial association

Local indicators of spatial association (LISA) are used to identify regional co-localization patterns (Anselin, 1995). After z-score standardization, spots are classified based on whether the primary feature exceeds the mean and whether the spatially lagged secondary feature is positive:

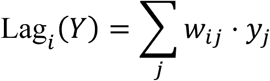

where *w_ij_* is the inverse Euclidean distance between spots *i* and *j*, globally normalized. This yields four spatial categories: high–high, low–low, high–low, and low–high associations.

#### Spatial ligand-receptor co-expression

Cell–ECM and ECM–ECM interaction pairs were obtained from MatriComDB (Lamba et al., 2025). For each ligand–receptor pair, a score is computed as:

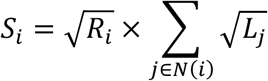

where *R_i_* is receptor expression at spot *i*, *L_j_* is ligand expression in neighboring spots, and *N*(*i*) denotes the six nearest neighbors in accordance with the Visium hexagonal grid. This formulation reflects the requirement for both receptor expression and local ligand availability. Enrichment within ECM niches is quantified using average log_2_ fold change:

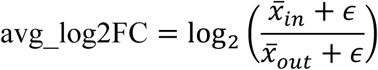

with *∈* = 10^−9^. Percentage differences capture changes in the proportion of active spots. Reciprocal interaction pairs are merged into bidirectional communication axes.

#### Differential matrisome gene expression analysis

In the R package version, MatriSpace allows for cross-sample differentially expressed matrisome gene (DEMG) analysis. DEMGs are identified using the Wilcoxon rank-sum test with Benjamini–Hochberg correction, filtering for genes expressed in at least 5% of spots in either group with a log2 fold-change threshold of 0.25. Significant genes are further assessed using a spatial mixed model (spaMM; Rousset & Ferdy, 2014) with Matérn covariance to distinguish spatially corrected from uncorrected results. For cross-sample comparisons, spot-level counts are aggregated into pseudo-bulk profiles and tested with DESeq2 (Love et al., 2014), controlling for composition cluster and sample effects. Results are annotated with matrisome division and category information.

### MatriSpace implementation

#### Software dependencies

MatriSpace was developed in R (4.3.3) using the Shiny framework (Chang et al., 2025) with bslib for interface components (Sievert et al., 2025). Core dependencies include Seurat (Hao et al., 2024), UCell (Andreatta and Carmona, 2021), HGNChelper (Oh et al., 2020), MERINGUE (Miller et al., 2021), and ScType (Nader et al., 2024). Visualization is implemented using *ggplot2* for static plots (Wickham, 2016), *plotly* for interactive graphics (Sievert, 2020), and *D3.js* for the spatial tissue viewer (Bostock et al., 2011).

#### Application maintenance

MatriSpace is part of the “MatriTool” ecosystem, which also features Matrisome AnalyzeR (Petrov et al., 2023) and MatriCom (Lamba et al., 2025), developed as part of the Matrisome Project (https://matrisome.org). As MatriTools serve as a central, actively curated framework for ECM-focused computational resources, MatriSpace will inherit continuous updates in matrisome annotations, gene classifications, and emerging biological insights. Likewise, novel functionalities will be added as progress in the ST field exposes or adopts novel analytical methods relevant to the scope of MatriSpace.

#### User support

MatriSpace provides contextual help via tooltips explaining each analysis parameter, step, and output. First-time users can access an interactive guided tour that walks them through the matrisome profiling and feature analysis workflows directly from the Home page. Additional support is available by contacting the development team at.

#### Case study and additional analytical methods

While all results presented in this manuscript were generated using MatriSpace, certain downstream analyses presented in the case study required extensions of the core analytical framework. In particular, transcription factors (TFs) regulating ECM genes were sourced from the TRRUST “Curated” database (http://www.grnpedia.org/trrust) (Han et al., 2018) and restricted to two non-overlapping sets, one mapping TFs to a peri-vascular ECM signature made from the combination of two signatures present in MatriSpace (hemostasis and peri-vascular ECM), and the other mapping TFs to the ECM-affiliated signature, and verifying their enrichment within core and border spots by inter-spot Pearson correlation. The Collagen-DDR signatures (*COL3A1-DDR1* and *COL1A1-DDR1*) were computed using the same robust scoring function as in MatriSpace, and their difference was used as a “dormancy score” based on the previously reported effects of the COL3A1-DDR1 interaction in supporting dormancy (Di Martino et al., 2022). The relationship between the two signatures and the expression of the proliferation marker *MKi67* was modeled using an additive generalized linear model (GLM) with a Gaussian link function, and the magnitude of partial (marginal) effects is reported. The spatial admixing of the BM and interstitial niches was tested with a custom cross-contiguity function computing, for all spots in group A, the average fraction of their k-nearest neighbors that belong to group B (i.e., a directed measure of how strongly A is spatially surrounded by B), the k-nearest neighbors’ structure being calculated using the *get.knn* function from the FNN package, setting k = 6. To facilitate interpretation, we classified each spot by whether its signature expression (per each of the niche signatures) lay above or below the median and assessed the spatial proximity of spots with dominant niche signals (one high and the other low) against 10⁴ random permutations of the same labels.

## RESULTS

### User interface of the online version of MatriSpace

#### MatriSpace input

The MatriSpace web application allows users to query a collection of open-access datasets or process their own datasets. The preloaded collection comprises 198 open-access datasets retrieved from various repositories (**Figure 1A**) and includes 18 datasets generated from healthy human tissue sections and 180 datasets generated from human tumor sections (**Figure 1B**), spanning most physiological systems. Once identified, users can load the dataset of interest via the Load sample button (**Figure 1C**). Alternatively, users can opt to analyze their own ST datasets, provided the file meets the stipulations listed in the Methods section. Namely, datasets should adhere to the current Seurat and SpatialExperiment standards (Seurat V5 and SpatialExperiment V3) and be saved as an RDS object. Internal MatriSpace functions will handle data preprocessing, if needed. Suitable objects are uploaded via the Load sample button (**Figure 1C**).

**Figure 1.**
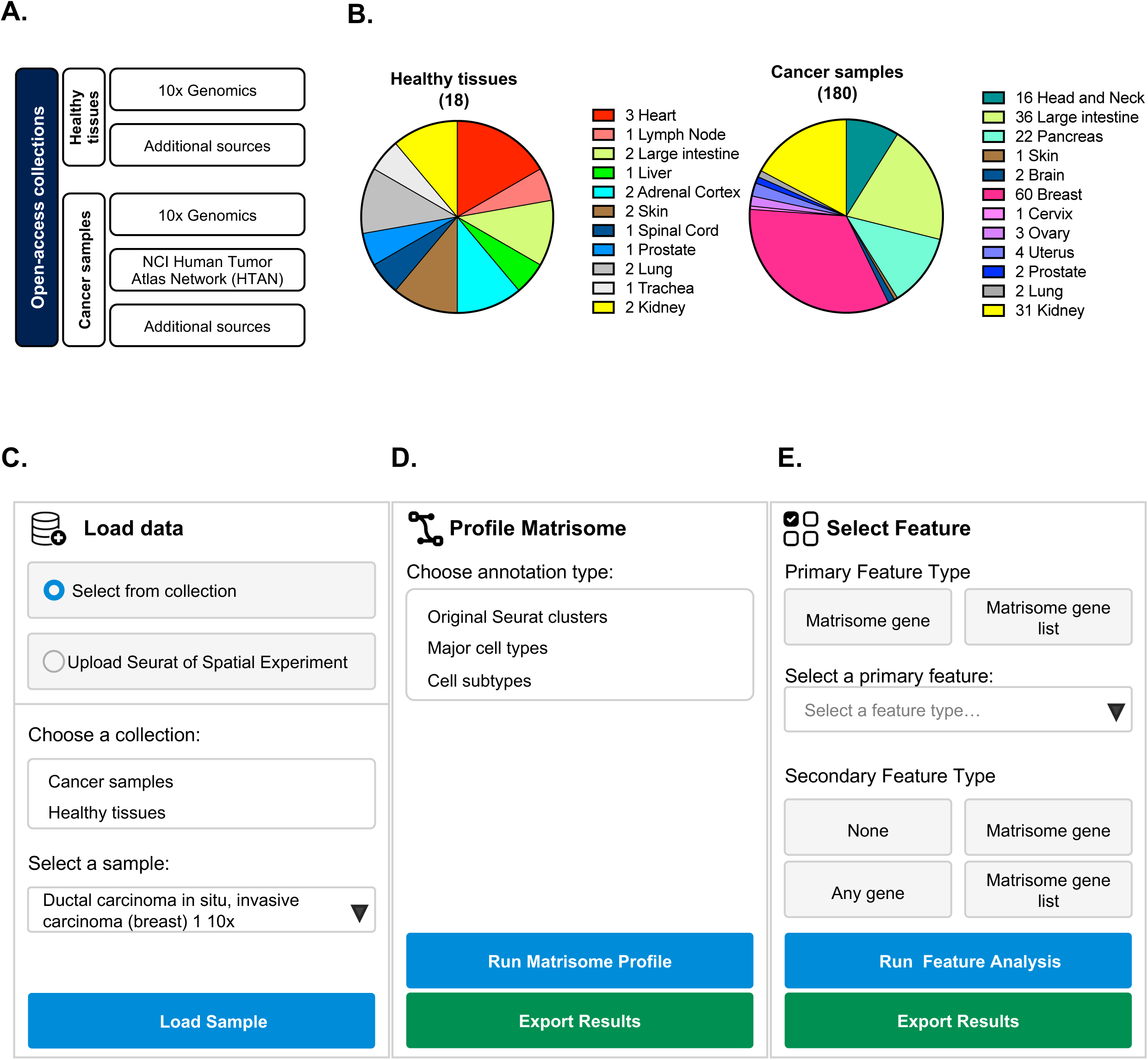
MatriSpace open-access dataset collections and query interface. **A.** Datasets were retrieved from public repositories, including 10x Genomics, the Human Tumor Atlas Network (HTAN), Gene Expression Omnibus (GEO), and Zenodo. **B. *Left:*** Pie chart represents the distribution of the 18 datasets of the healthy tissue collection by tissue of origin. ***Right:*** Pie chart represents the distribution of the 180 datasets of the cancer sample collection by tissue of origin. *See also Table S1*. **C.** Data input panel: Users can select a dataset from the MatriSpace collection or upload their own processed objects. **D.** Matrisome profiling analysis panel: Users can select a cell annotation type to compute the matrisome profile (blue button) and export result files (green button). **E.** Feature analysis panel: Users can interrogate specific features using a two-level selection system. Upon feature selection, users can perform feature analysis (blue button) and export result files (green button).

#### Query parameters

Upon loading a dataset, users must select an annotation type from the metadata layer of the data in the Profile matrisome panel. This will stratify all downstream analyses by a biological layer of interest, *e.g.,* cell type labels, clusters, or histologically or pathologically defined regions. Note that, for the open-access data provided with the online version of MatriSpace, cell annotations from a harmonized deconvolution pipeline (*see Methods*) are provided. From that same panel, users can then opt to Run matrisome profile to obtain a global view of matrisome gene sets distribution in the sample of interest with spatial resolution (**Figure 1D**). Alternatively, users can perform a more focused analysis using the Select feature panel (**Figure 1E**). When opting to run a feature analysis, users can interrogate the expression of a single matrisome gene or of a matrisome gene set. Users can also perform co-analysis of two features, the secondary feature including, in addition to matrisome genes and gene sets, any genes (*see below*).

### MatriSpace output

To illustrate the functionalities of MatriSpace, we will use the open-access 10x Genomics breast ductal carcinoma *in situ* with invasive carcinoma dataset (Janesick et al., 2023) processed through the online MatriSpace application with the “major cell type” as the annotation mode throughout this manuscript.

#### Dataset visualization

Upon loading the dataset, the hematoxylin and eosin (H&E)-stained tissue section (**Figure 2A, left panel**) and the corresponding cluster map depicting the spots profiled and the nature of the cell clusters present in this sample will appear in the main window (**Figure 2A, right panel**). Users are also provided with a set of metrics in cards at the top of the page, including the number of sampling spots collected (here, 4,992) and the number of genes and matrisome genes represented in the dataset (here, 18,045 and 948, respectively). Clicking on the Learn more about this sample button will take users to the link of the original dataset in a separate window (here, https://cf.10xgenomics.com/samples/spatial-exp/2.0.0/CytAssist_FFPE_Human_Breast_Cancer/CytAssist_FFPE_Human_Breast_Cancer_we <u>b_summary.html</u>).

**Figure 2.**
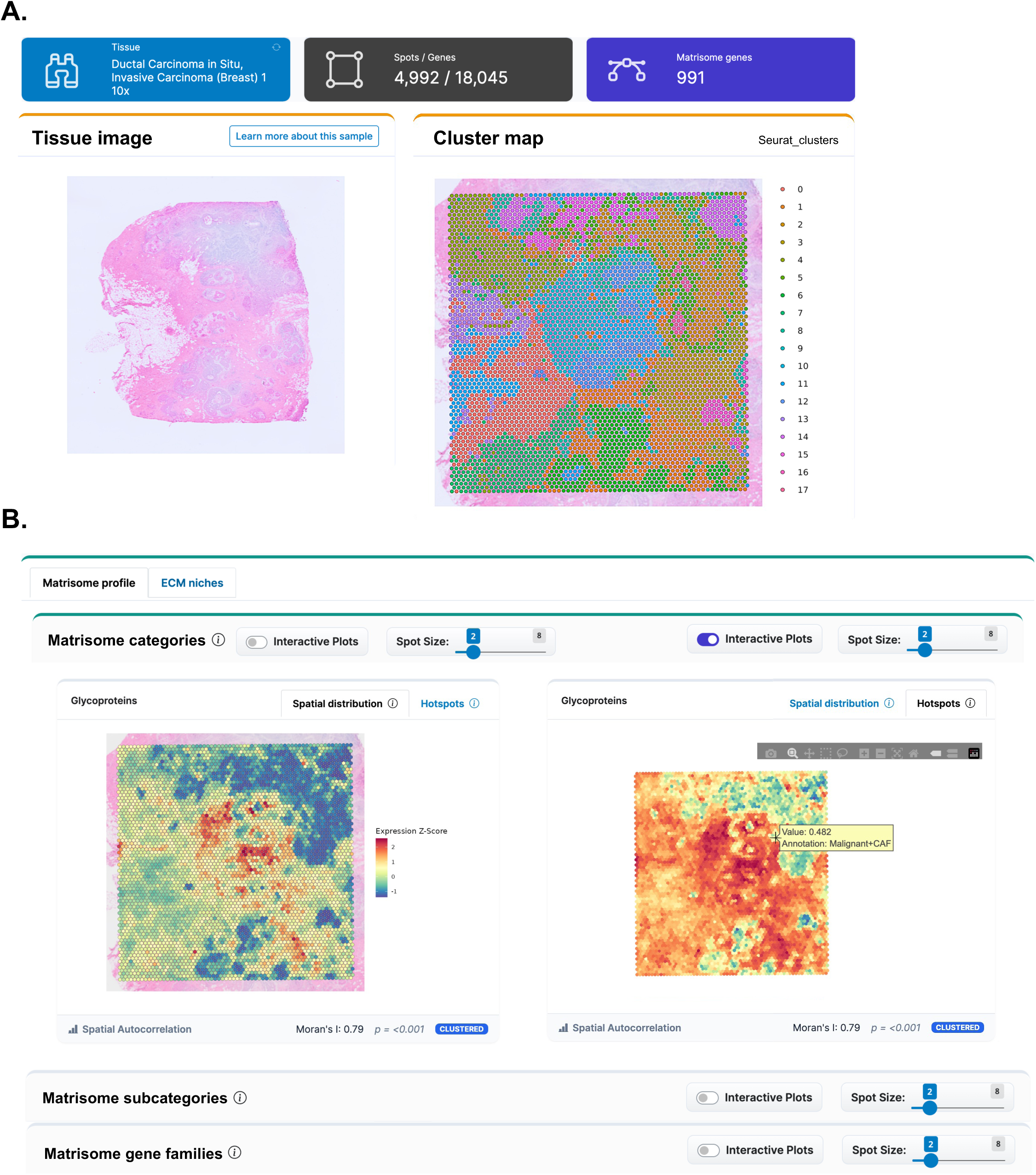
MatriSpace cell type annotation and matrisome profiling output. We illustrate the functionalities of MatriSpace in Figures 2 to 5 using the following BrK dataset from https://cf.10xgenomics.com/samples/spatial-exp/2.0.0/CytAssist_FFPE_Human_Breast_Cancer/CytAssist_FFPE_Human_Breast_Cancer_web_summary.html (Janesick et al., 2023). **A.** Hematoxylin and eosin (H&E) staining conducted pre-CytAssist is shown alongside the Visium spatial plot with default Seurat-derived cell type clusters. **B.** Spatial visualization of matrisome signature expression, illustrated here by ECM glycoproteins, is shown as either a spatial distribution (*left*) or a hotspot map (*right*). Users can switch to an interactive plot and/or adjust spot size. Within the interactive plot, users can zoom into regions of interest, select specific areas, or download the image using the interactive panel. Hovering over individual spots displays expression value and annotated cell type.

#### Matrisome profiling

After selecting a dataset, users can choose the cell type annotation they wish to use and run an analysis of the expression of all matrisome genes quantified in the dataset by selecting the Run Matrisome Profile button (**Figure 1B**). This analysis will return a set of expression maps grouped into two sets: matrisome expression profile maps and maps representing the spatial distribution of cells expressing gene panels characteristic of defined functional ECM niches.

The Matrisome profile tab displays spatial distribution and hotspot maps for a panel of matrisome gene sets. The top panel includes maps for each of the six canonical matrisome gene categories encoding, respectively, ECM glycoproteins, collagens, proteoglycans, ECM-affiliated proteins, ECM regulators, and other secreted factors defined based on protein domain-based organization (Hynes and Naba, 2012; Naba et al., 2012). The middle panel includes maps showing the expression of matrisome genes classified into four functional subcategories: 1) genes encoding proteins found in the perivascular region, 2) genes encoding ECM proteins involved in hemostasis, 3) genes encoding proteins participating in the formation of elastic fibers, and 4) genes encoding proteins with growth-factor binding activities. The bottom panel includes maps reporting the expression of four matrisome gene families: the laminins, which are glycoproteins of the basement membrane ECM, matricellular proteins that are glycoproteins that associate with fibrillar ECM proteins, and syndecans and glypicans that are transmembrane proteoglycans (all gene sets are available in **Supplemental Table S2**). In combination with the identification of histological structures from the H&E image, and the nature and distribution of the different cell populations across the tissue section (cluster map), these matrisome maps can inform users on the spatial organization of the populations expressing subsets of matrisome genes in context (*see examples below*).

Users can access different maps for each gene set: a spatial distribution map and a hotspot map, as shown in **Figure 2B** for the matrisome glycoprotein gene set. The spatial distribution map (relative expression) uses a Z-score to represent how much a spot’s expression deviates from the tissue’s median expression level. This type of map can be used to identify regions where the expression of a given matrisome gene set is statistically up- or down-regulated as compared to the tissue’s baseline. In contrast, the hotspot map (absolute expression) uses a log-transformed score to show the full expression range of the selected matrisome gene set across the tissue section. Users can explore this type of map to identify hotspots of expression, interpreted as the main cellular source(s) of matrisome proteins encoded by a given gene set. All maps are accompanied by spatial autocorrelation statistics (Moran’s I) indicative of whether data patterns are clustered, dispersed, or randomly distributed, by location and value. Note that all the maps can be made interactive by toggling the Interactive Plots button located in the upper-right corner. Activating this functionality will allow users to download maps as .png images, zoom in or out on specific regions, and obtain expression data for a specific spot by hovering over it.

#### Identification of functional ECM niches

Under the ECM niches tab, users will find a set of plots representing the expression of genes encoding proteins that form distinct functional ECM niches, namely the basement membrane ECM and the interstitial ECM (**Figure 3A)**. This tab also features ligand–receptor co-expression analyses identifying enriched ECM communication axes associated with selected niches. Specifically, the ECM niche annotation map is an interactive map that classifies each spot into a dominant ECM niche based on marker gene expression (**Supplemental Table S2D)**. This representation provides a functional, high-level annotation of a tissue’s microenvironment. This map is interactive, and users can set up visualization parameters such as spot size and spot opacity. The niche score plots represent these signals as continuous variables capturing gradients in the expression of genes that are part of defined niches and revealing spatial transitions. The Niche distribution panel includes a bar graph that summarizes the average expression levels of the gene signature of each of the two ECM niches, for each cell type (**Figure 3B**). It thus allows the identification of the cell type(s) or tissue regions that are the primary contributors to the interstitial or basement membrane ECM gene expression programs.

**Figure 3.**
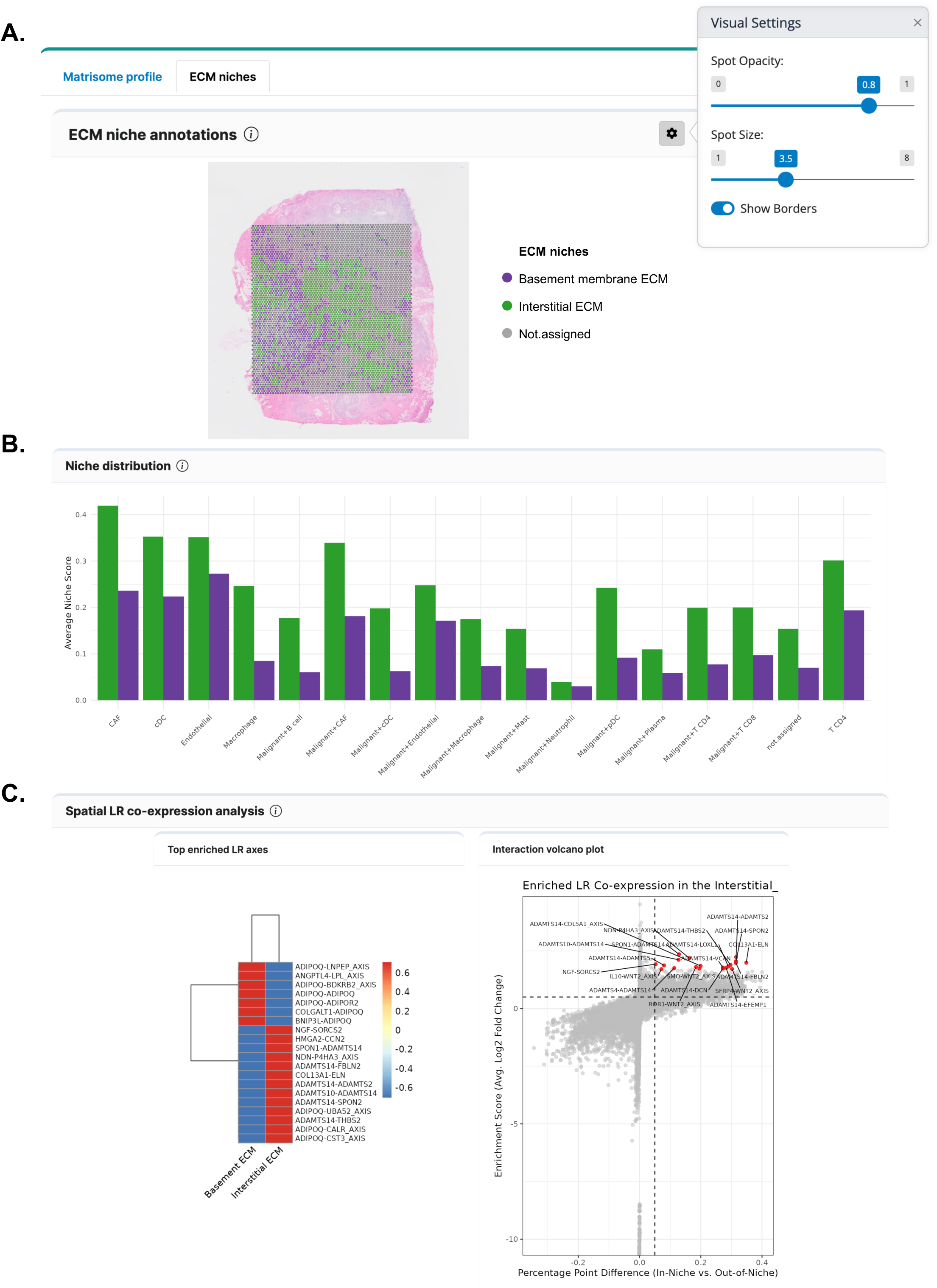
Identifying the localization and patterning of ECM niches with MatriSpace. **A.** Upon selecting the “Niche output” feature, a Visium spatial plot overlaid with H&E staining is displayed, showing the spatial distribution of ECM niches (*i.e.*, basement membrane ECM or interstitial ECM) assigned to individual spots. Users can adjust the spot size and transparency using the visual setting button (right top corner). **B.** Histogram displays the average niche score for each annotated cell cluster. Color codes represent the basement membrane ECM (purple) and the interstitial ECM (green) niches. **C.** Heatmap and volcano plot visualize the analysis scores of the spatial ligand-receptor pairs within each ECM niche. S*ee also Methods*.

Last, the Spatial LR co-expression analysis panel returns the identification of potential signaling hubs by measuring the co-expression of a ligand (L) in one spot and its corresponding receptor(s) (R) in neighboring spots. The output of this analysis is a heatmap that displays the top enriched ligand-receptor pairs or “LR axes” across annotated niches (**Figure 3C, left panel**), providing a high-level overview of the most active communication patterns. Users can also visualize the data as a volcano plot depicting LR pairs for a single selected niche, highlighting those that are most significantly enriched in that specific environment (**Figure 3C, right panel**). Users should note that this analysis can be computationally intensive and may take several minutes to complete, though extensive optimization generally confines longer runtimes to only the most extreme cases. The other functionalities of the MatriSpace will thus be on hold while LR co-expressions are computed.

#### Feature query selection and output

After selecting a dataset, users can also opt to run a custom query by expanding the Select Feature query panel (**Figure 1E**). As an example, we performed a feature analysis using the matrisome genes *COL1A1* (primary feature) and *COL1A2* (secondary feature). These genes encode respectively the α1 and α2 chains of collagen I, which assemble to form a functional collagen I trimer composed of two α1 and one α2 chains (Ricard-Blum, 2011).

As presented above, the Tissue Overview Map orients users to the identity and distribution of the different cell populations present in the tissue section of interest (**Figure 4A**). The Feature Expression box-and-whisker plots depict the expression levels of *COL1A1* and *COL1A2* across the different cell populations (**Figure 4B**). As expected, we found that both genes are predominantly expressed in cancer-associated fibroblasts (CAFs) and that *COL1A1* is expressed at a higher level than *COL1A2*, likely accounting for the stoichiometry in which their gene products contribute to the functional trimeric protein. The Feature maps, overlaid onto the tissue overview, represent the spatial distribution and expression levels of the two selected features (**Figure 4C**); while the Co-expression map depicts areas where the two features co-localize, are mutually exclusive, or are jointly absent (**Figure 4D**). Here, as expected, we found that at the whole-tissue section level, the expression of *COL1A1* and *COL1A2* appears to be highly correlated (Pearson coefficient: r = 0.91 and p<0.001; spatial cross-correlation: 0.76). Last, the LISA map represents regional co-localization patterns with a complementary spatial approach, particularly useful when patterns of co-occurrence are discontinuous or non-linear, for example, between ligands and receptors; here, most of the co-expression occurs along the borders of the area otherwise marked by high *COL1A1* expression (**Figure 4E**).

**Figure 4.**
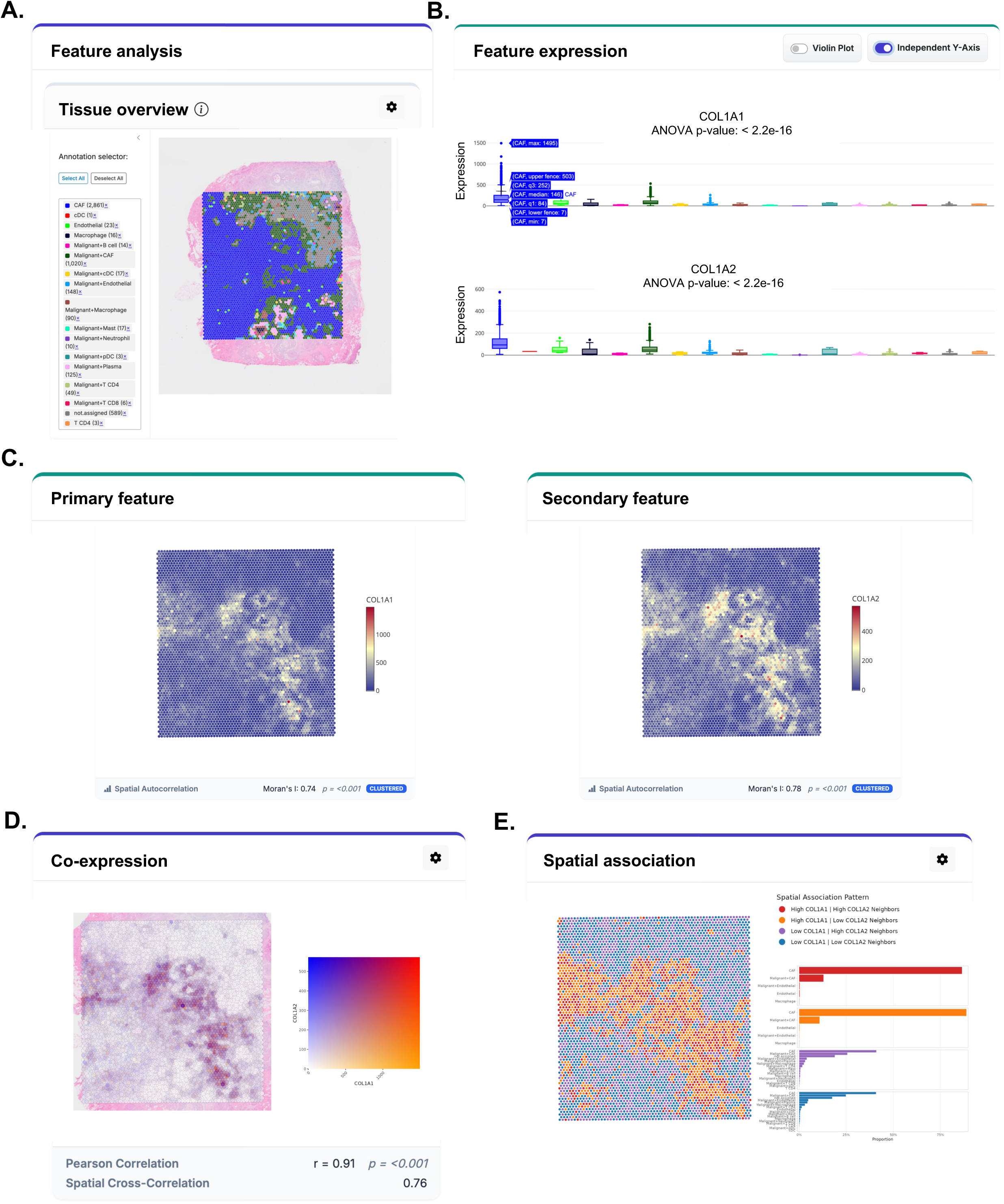
MatriSpace feature analysis output. **A.** Annotated cell types are displayed on the Visium spatial plot overlaid with H&E staining, with colors indicating cell-type identity. Users can use the annotation selector panel to filter cell type(s) of interest. **B.** Boxplots show the expression of the selected primary and secondary features, here *COL1A1* and *COL1A2*, across cell types; hovering over individual boxes reveals the cell type identity and quartile values. Users can switch to a violin plot or enable an independent Y-axis. **C.** Visium spatial plots depict the spatial distribution of cells expressing *COL1A1* (*left*) and *COL1A2* (*right*). **D.** Co-expression analysis of *COL1A1* and *COL1A2*. The color of each spot indicates the expression patterns: orange for high *COL1A1* expression, blue for high *COL1A2* expression, and red for high co-expression. **E.** Local indicators of spatial association (LISA) score of selected features are displayed on the Visium plot. The color of each spot represents one of the following association patterns: red (high expression of both primary and secondary features), orange (high expression of primary feature), purple (high expression of secondary feature), and blue (low expression of both primary and secondary features). The corresponding histograms show the distribution of cell types across spatial association patterns.

#### Result export

Upon running a Matrisome Profile or a Feature analysis, users can export the results (compressed .zip folder or individual files) using buttons located in the control panels (**Figures 1D and 1E**). Results files include a README.txt file that describes the content of each data and image file, a Data folder, and a Plot folder. The Data folder will contain a .rds file (here, ‘Breast_cancer_Ductal_Carcinoma_In_Situ_Invasive_MatriSpace.rds’) of the complete Seurat object with all analysis results, a ‘spot_metadata.csv’ file listing the metadata (*i.e.*, coordinates, annotations, scores) for each spot profiled, and a ‘matrisome_scores.csv’ file that compiles all calculated matrisome gene signature scores for each spot. If users have further performed a feature analysis, that Data folder will also contain additional files, including a ‘main_feature_data.csv’ file listing the expression values for the selected features for each spot, a ‘spatial_statistics_summary.csv’ file providing a summary of all calculated correlation statistics, an ‘lr_coexpression_stats.csv’ file that compiles ligand-receptor co-expression enrichment statistics per niche and an ‘lr_coexpression_means.csv’ compiling the mean co-expression scores for each ligand-receptor axis, per ECM niche. The Plot folder contains all generated plots, including distribution maps and hotspot maps, in two formats: .png and .pdf. The .png files under the ‘PNG_for_Viewing’ subfolder provide high-resolution .png files, while the .pdf files are vector-based and suitable for publication and editing. Note that the export can take several minutes, depending on the dataset size.

### Dissecting the microenvironmental heterogeneity of breast cancers with MatriSpace

To illustrate the usability of MatriSpace, we performed a comprehensive analysis of a recently published breast cancer (BC) spatial transcriptomic dataset generated using the Visium Cytassist platform (Janesick et al., 2023). This sample provides a high-resolution view of BC histopathology and disease architecture, as it presents both ductal carcinoma *in situ* (DCIS) and invasive ductal carcinoma (IDC) regions surrounding a central stromal/immune compartment enriched for transcriptionally diverse cell populations (**Figure 5A**). This sample also shows areas with high and low cellularity (see #RNA count panel) and areas with higher or lower gene expression (see #RNA features panel). At the section level, gene signature scores corresponding to the interstitial (I) and basement membrane (BM) ECMs are concentrated within the stromal regions (**Figure 5B**). Notably, the two niche scores display substantial co-localization, yielding a significant positive correlation (R = 0.7, p < 0.001; **Figure 5C**). This might indicate that the same cell population contributes to the expression of genes encoding proteins of the interstitial and basement membrane ECMs, warranting further investigation. This may also simply reflect the fact that the spatial resolution of the dataset (∼50 µm spot diameter) limits our ability to resolve fine-grained expression patterns and distinguish signals from adjacent but biologically distinct microenvironments. Importantly, however, this global correlation does not completely mask spatial heterogeneity, as BM and I scores do not overlap uniformly. Regions dominated by one program (*i.e.*, high BM/low I or high I/low BM) show markedly reduced spatial proximity (cross-contiguity = 0.08, p < 0.001; **Figures 5C and 5D**) compared to areas where both signals are either high or low. Together, these findings support the concept that BM and I niches are not independent but co-activated components of a dynamic expression pattern continuum, whose spatial decoupling (or overlap) captures distinct ECM gene expression programs within the tumor microenvironment (Hunter et al., 2021; Winkler et al., 2020). In addition to these observations, we found that the distribution of dominant ECM niche signals varied across histopathological compartments and tumor states (**Figure 5E**). Specifically, DCIS regions exhibit higher basement membrane and interstitial niche scores than IDC areas, which show reduced matrisome gene expression. This is consistent with evidence that pre-invasive lesions retain structured ECM organization, whereas invasive tumors undergo basement membrane disruption and matrix remodeling, relying less on coordinated ECM gene expression and more on stromal reprogramming. In contrast, intact tissue compartments, including *FABP4*-positive adipose and *KRT17*-positive myoepithelial regions, display selectively elevated BM scores that may indicate the retention of intact epithelial and adipose gene programs in these areas. Of note, adipocytes present the particularity of being surrounded by a specialized basement membrane. Together, these patterns illustrated how MatriSpace can resolve biologically meaningful, spatially constrained ECM gene expression programs that track with cell content, tissue organization, and tumor progression.

**Figure 5.**
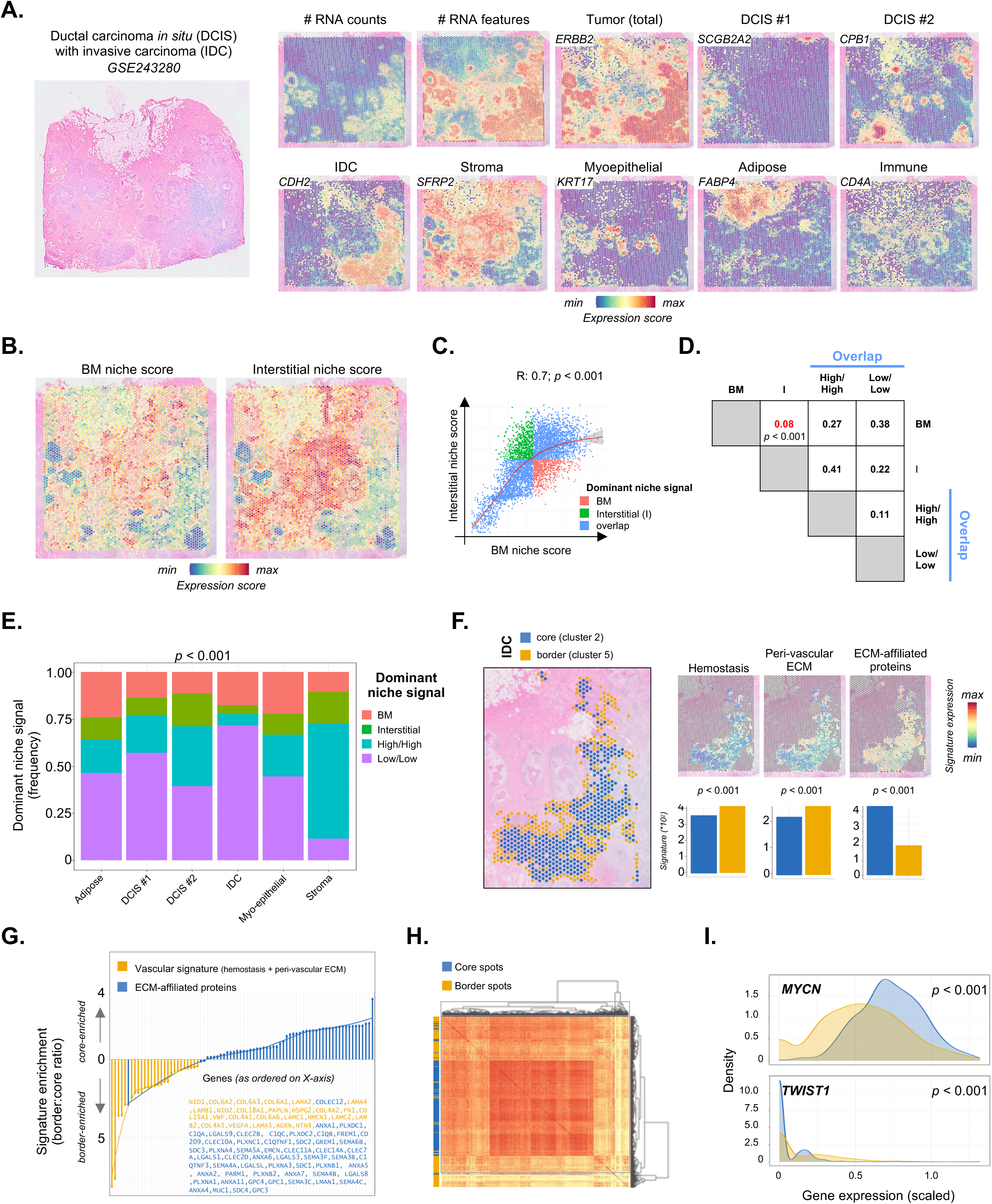
Deciphering breast cancer microenvironmental heterogeneity with MatriSpace. **A.** High-resolution hematoxylin & eosin staining (*left*) and broad cell types as per the original publication (GSE243280; *right*). DCIS: ductal carcinoma *in situ,* IDC: invasive ductal carcinoma. Of note, the panels in Figure 5 are all rotated by 90 degrees relative to the images in Figures 2 to 4 (which used the orientation registered as metadata in the Visium dataset) to match the images in the original study by Janesick *et al*. **B.** Visium spatial plots depicting expression scores for the basement membrane (BM) and the interstitial ECM niches. **C.** Correlation plot (and Pearson’s coefficient, R) illustrates the relation between the interstitial and basement membrane (BM) ECM niche scores. Note how spots with a dominant niche signal (where one signature is above the mean and the other below) segregate to the opposite ends of the distribution. **D.** Contiguity analysis of the niche types. The p-value resulting from 1*10^4^ random-label permutations was calculated to assess statistical significance. Note the significant spatial separation between spots where one niche score dominates. *See also Methods*. **E.** Bar graph represents the frequency of each niche by broad cell types. The p*-*value from a Chi-square test with continuity correction was calculated to assess statistical differences. **F.** Definition of border and core IDC regions based on unsupervised Seurat clustering *(left)* and differential expression of ECM signatures between the regions. The p*-*value was calculated using the Wilcox test to assess statistical differences. **G.** Enrichment (border *vs.* core) values for the genes in the signatures identified in (**F**). For clarity, genes belonging to the “hemostasis” and “peri-vascular ECM” signatures were collapsed into a single “vascular” signature. **H.** Expression of transcription factors potentially regulating the vascular and ECM-affiliated proteins signature along IDC border and core spots. *See also Methods*. **I.** Examples of differentially regulated transcription factors from those identified in (H).

For higher-resolution analyses, MatriSpace can compute robust spatial enrichment statistics for structural and functional matrisome signatures, which can be readily integrated with associated metadata to resolve transcriptional programs at discrete regional locations. Leveraging the unsupervised clusters generated through the standard spatial Seurat workflow (Hao et al., 2024), we annotated core and border regions within the invasive ductal carcinoma (IDC) lesion and interrogated compartment-specific ECM programs (**Figure 5F**). Strikingly, these regions exhibited a strong polarization toward distinct ECM states. Namely, border spots were significantly enriched for genes encoding matrisome components involved in hemostasis and the formation of the peri-vascular niche; in contrast, core spots showed marked enrichment for ECM-affiliated proteins. Differential expression analysis confirmed that this segregation was not driven by isolated markers but rather reflected coordinated upregulation of multiple genes within each signature (**Figure 5G**), consistent with transcriptional specialization. To contextualize these differences, we applied the same strategy previously implemented in MatriCom (Lamba et al., 2025) and interrogated the TRRUST database (Han et al., 2018) to identify transcription factors (TFs) reported to regulate genes within the vascular and ECM-affiliated programs. This analysis revealed a clear bifurcation of core and border spots along transcriptional regulatory axes, forming distinct “blocks” of coordinated expression of the differential TFs (**Figure 5H**). At the ends of this spectrum, *MYCN* was preferentially enriched in core regions, whereas *TWIST1* predominated at the border (**Figure 5I**). This distribution aligns with established roles of MYCN in proliferative and metabolically active tumor states and of TWIST1 in mesenchymal transition, invasion, and vascular remodeling (Qin et al., 2026), supporting the biological relevance of the spatially polarized ECM programs identified.

### MatriSpace enables the identification of ECM-dependent signaling pathways with spatial context

At maximal granularity, MatriSpace enables spatial resolution of individual genes or gene pairs, provided at least one component belongs to the matrisome. This flexibility allows MatriSpace outputs to be seamlessly integrated with complementary analytical frameworks, extending its applicability beyond the native functionalities of the application and supporting downstream analyses in external pipelines. To illustrate this usability and pinpoint discrete changes in ECM gene expression events at the locoregional level, we focused on a small intra-adipose tumor lesion originally classified as triple-positive (*ERBB2⁺, ESR1⁺, PGR⁺*) DCIS (**Figure 6A**), in contrast to the predominantly double-positive (*ERBB2⁺, ESR1⁺*) phenotype of the surrounding tumor (Janesick et al., 2023). This lesion exhibited strong depletion of *MKI67*, along with strong expression of *DDR1* and *COL3A1*, and a reduction in *COL1A1* (**Figure 6A**). We used the expression values of these genes to construct a *COL3A1/DDR1* and a *COL1A1/DDR1* signature using the same formulation as in MatriSpace and defined a “dormancy score” as their difference. Results show a relative enrichment of the LR pair associated with dormancy (*COL3A1*/*DDR1*) associated with low *MKI67* expression, suggesting dormancy within this tumor area (**Figure 6B**). To more formally test this hypothesis, we then fitted a generalized linear model (GLM) to relate each signature to proliferation, allowing the effect of one signature to be estimated while controlling for the other. We found that the *COL1A1*-based score showed a strong positive association with proliferation, whereas the *COL3A1*-based score showed a significantly negative association (**Figure 6C**). Together, these findings support the notion that this discrete triple-positive lesion occupies a distinct collagen III–defined ECM niche that biases DDR1 signaling toward dormancy rather than expansion, distinguishing it from the proliferative collagen I–rich regions that characterize the remainder of the tumor.

**Figure 6.**
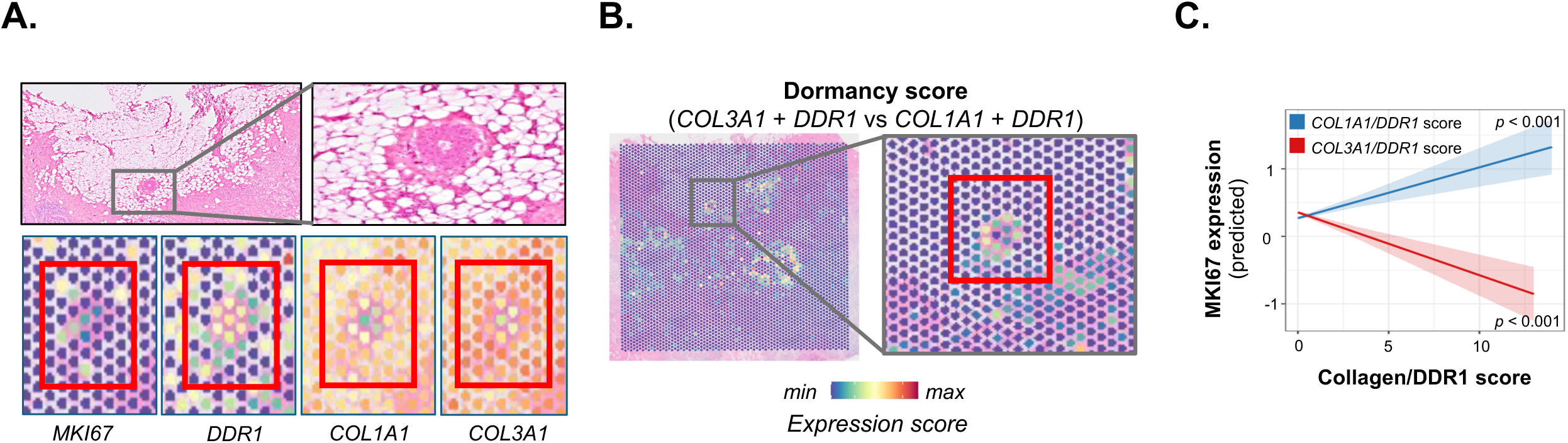
Identifying ECM-dependent signaling pathways and functional niches with spatial context. **A.** Identification of a triple-positive intra-adipose DCIS#2 lesion as from the original publication, and expression of the proliferation marker *MKI67*, the fibrillar collagens *COL1A1* and C*OL3A1*, and the discoidin domain receptor 1 (*DDR1*). **B.** Relative enrichment of an ECM ligand/receptor dormancy score (*COL3A1* + *DDR1* score *vs. COL1A1* + *DDR1*) coupled to low *MKI67* expression suggests dormancy within this tumor area. **C.** Generalized linear model of *MKI67* expression against *COL3A1* + *DDR1* and *COL1A1* + *DDR1* expression scores shows a positive correlation between the *COL1A1*/*DDR1* signature and *MKI67*expression independently of the *COL3A1*/*DDR1* (p < 0.001), and a negative correlation between the *COL3A1*/*DDR1* signature and *MKI67*expression (p < 0.001).

## DISCUSSION AND CONCLUSION

MatriSpace offers a conceptual and methodological advance in the field of spatial transcriptomic dataset analysis by explicitly focusing on modeling the ECM at multiple levels of organization rather than treating matrisome genes as downstream markers of cell populations. Unlike existing tools that primarily focus on cell-type deconvolution, clustering, or generic pathway enrichment, As illustrated in this manuscript, MatriSpace reframes ECM biology as a spatially structured system of interacting programs, enabling direct interrogation of tissue-wide niche states as well as more locoregionally-defined gene signatures and interactions.

A key advance of MatriSpace lies in its ability to infer functional ECM axes at high spatial granularity, including gene-level and gene-pair–level interactions constrained by matrisome membership. This enables analyses that go beyond conventional module scoring by capturing directional and relational structure between ECM components and other biological programs. Importantly, this framework does not merely quantify expression differences but provides means to dissect spatial transitions between ECM states, key to developmental and pathological processes, and to uncover biologically meaningful structures that are often obscured in standard cell-centric analyses.

Despite presenting advantages over existing pipelines, several limitations remain. A fundamental constraint is imposed by the resolution of current sequencing-based spatial transcriptomics approaches, where each spot (∼50 µm) aggregates transcripts from multiple cells. As a consequence, observed co-localization patterns may reflect true biological coupling or the averaging of adjacent microenvironments with distinct ECM gene expression levels, potentially contributing to different ECM states. This limitation is particularly relevant when interpreting gradients or transitions between basement membrane–rich and interstitial matrix–rich regions. Fortunately, technological and methodological advances in the field (*e.g.*, the introduction of “HD” platforms with significantly more granular outputs) as well as the expansion of capture panels for *in-situ*-based methods to include more ECM signals will likely change this outlook in the near future. We anticipate being able to update MatriSpace towards these standards in the coming years.

With now well-established discrepancies between gene expression levels and protein abundance (Liu et al., 2016), a second limitation concerns the reliance on transcript abundance as a proxy for ECM structure and function or cell-ECM interactions (Lamba et al., 2025). This is further exacerbated by the fact that the ECM is a heavily post-transcriptionally regulated system, where protein deposition, crosslinking, degradation, and mechanical organization can only be partially inferred from mRNA levels. Thus, while MatriSpace captures transcriptional programs associated with ECM remodeling, its output should be interpreted with caution, particularly in contexts where post-translational regulation dominates ECM behaviors. However, we envision that the development of spatially resolved proteomic technologies will overcome this critical limitation (Bodenmiller, 2024; Fan, 2024; Nordmann et al., 2024), and contribute to accelerating ECM research. And accordingly, we will aim to support this by further developing MatriSpace to be compatible with future dataset formats.

In conclusion, MatriSpace provides a framework that focuses on ECM biology as a spatially explicit analytical system, enabling the study of ECM organization as a continuum of interacting gene expression programs within tissue microenvironments. While current technological and biological constraints limit the resolution and functional inference of ECM states, the approach establishes a foundation for integrating spatial transcriptomics with mechanistic models of tissue architecture and opens future avenues for multi-modal extensions incorporating proteomic and imaging-based ECM readouts.

## TOOL AND DATA AVAILABILITY

- The web-based MatriSpace is deployed as a Shiny Application and is freely accessible at https://matrinet.shinyapps.io/matrispace/
- The locally deployable R Shiny application is available at https://github.com/izzilab/matrispace-app
- The MatriSpace R package is available at https://github.com/izzilab/matrispace

## ACKNOWLEDGMENTS

The authors would like to thank the members of the Izzi and Naba laboratories for their feedback on MatriSpace and this manuscript.

## AUTHOR CONTRIBUTIONS

Conceptualization: VI, AN

Methodology: AO, VI, AN

Software: AO, PP, VI

Data curation: AO, DC, VI, AN

Investigation: AO, DC, VI, AN

Visualization: AO, DC, VI, AN

Funding acquisition: VI, AN

Project administration and Supervision: VI, AN

Writing – original draft: AO, DC, VI, AN

